# Reduced Successor Representation Potentially Interferes with Cessation of Habitual Reward-Seeking

**DOI:** 10.1101/2020.09.15.297655

**Authors:** Kanji Shimomura, Ayaka Kato, Kenji Morita

## Abstract

Difficulty in cessation of drinking, smoking, or gambling has been widely recognized. Conventional theories proposed relative dominance of habitual over goal-directed control, but human studies have not convincingly supported them. Referring to the recently suggested “successor representation” of states that enables partially goal-directed control, we propose a dopamine-related mechanism potentially underlying the difficulty in resisting habitual reward-seeking, common to substance and non-substance reward. Consider that a person has long been taking a series of actions leading to a certain reward without resisting temptation. Given the suggestions of the successor representation and the dimension reduction in the brain, we assumed that the person has acquired a dimension-reduced successor representation of states based on the goal state under the established non-resistant policy. Then, we show that if the person changes the policy to resist temptation, a large positive reward prediction error (RPE) becomes generated upon eventually reaching the goal, and it sustains given that the acquired state representation is so rigid that it does not change. Inspired by the anatomically suggested spiral striatum-midbrain circuit and the theoretically proposed spiraling accumulation of RPE bias in addiction, we further simulated the influence of RPEs generated in the goal-based representation system on another system representing individual actions. We then found that such an influence could potentially enhance the propensity of non-resistant choice. These results suggest that the inaccurate value estimation in the reduced successor representation system and its influence through the spiral striatum-midbrain circuit might contribute to the difficulty in cessation of habitual reward-seeking.

## Introduction

Difficulty in cessation of drinking, smoking, gambling, or gaming, even with strong intention, has been widely recognized. Reasons for this, and whether there are reasons common to substance and non-substance reward, remain elusive. Although much effort has been devoted to developing clinical programs including technology-based therapies (e.g. (Gustafson *et al.*, 2014; Kato *et al.*, 2020b); reviewed in (Newman *et al.*, 2011; Haskins *et al.*, 2017)), the lack of mechanistic understanding of the undesired habit is an obstacle for further improvement. Computational modeling has become a powerful approach to elucidating the mechanisms of psychiatric disorders including addiction (Montague *et al.*, 2012; Wang & Krystal, 2014; Huys *et al.*, 2016; Kato *et al.*, 2020a). However, it appears that relatively less focus has been given to non-substance, compared to substance, addiction, although there have been proposals (e.g., (Redish *et al.*, 2007; Piray *et al.*, 2010; Ognibene *et al.*, 2019)). Also, previous computational studies appear to typically focus on the difficulty in withdrawal from severe addiction, including the problems of relapse. However, presumably a larger population of people have established milder, stable habitual reward-seeking behavior for years but then decide to quit such behavior because it potentially causes health or socioeconomic problems rather than because the behavior itself is immediately problematic as pathological addiction. In the present study, we explored possible computational and neural circuit mechanisms for the difficulty in resisting such stably established habitual behavior to obtain reward, with the following four streams of findings and suggestions in mind:

1. ***Involvement of the dopamine (DA) system in both substance and non-substance addiction*** The DA system has been suggested to be crucially involved in substance addiction (Berke & Hyman, 2000), possibly through drug-induced DA acting as a fictitious RPE that cannot be canceled out by predictions (Redish, 2004; Keiflin & Janak, 2015). However, there have also been implications of possible involvements of the DA system in non-substance addiction (Grant *et al.*, 2010). Specifically, possible relations of medicines of Parkinson disease to pathological gambling (Dodd *et al.*, 2005; Voon *et al.*, 2006), as well as similar changes in the DA system in addiction to substance and non-substance such as game (Thalemann *et al.*, 2007) or internet (Hou *et al.*, 2012), have been suggested.
2. ***Goal-directed and habitual behavior and their neural substrates, and their relations to addiction*** It has been suggested that there are two behavioral processes, namely, goal-directed and habitual behavior, which are sensitive or insensitive to changes in outcome values and/or action-outcome contingencies, respectively (Balleine & Dickinson, 1998; Balleine & O’Doherty, 2010; Dolan & Dayan, 2013). They are suggested to be hosted by distinct corticostriatal circuits, specifically, those including ventral/dorsomedial striatum (or caudate) and those including dorsolateral striatum (or putamen), respectively (Corbit *et al.*, 2001; Yin *et al.*, 2004; Yin *et al.*, 2005), where ventral-to-dorsal spiral influences have been anatomically suggested (Haber *et al.*, 2000; Joel & Weiner, 2000). Computationally, goal-directed and habitual behavior have been suggested to correspond to model-based reinforcement learning (RL) and model-free RL, respectively ((Daw *et al.*, 2005); but see (Dezfouli & Balleine, 2012) for a critique of model-free RL as a model of habitual behavior). It has been suggested that addiction can be caused by impaired goal-directed and/or excessive habitual control (Everitt & Robbins, 2005; 2016). This is supported by multitudes of animal experiments, and there also exist findings in humans in line with this (Gillan *et al.*, 2016). However, it has also been shown that human addicts often show goal-directed behavior, such as those sensitive to outcome devaluation (Hogarth *et al.*, 2019), although there are mixed results (as reviewed in (Hogarth *et al.*, 2019)) and sensitivity can also differ between appetitive and aversive outcomes as shown for cocaine addiction (Ersche *et al.*, 2016). Also, there have been proposals of many different possible causes for addiction, including those related to each of the two control systems and/or in their interactions, the way of state representation, or the hierarchical organization of the learning system (e.g., (Redish *et al.*, 2007; Redish *et al.*, 2008; Keramati & Gutkin, 2013; Keramati *et al.*, 2017; Ognibene *et al.*, 2019)).
3. ***Intermediate of goal-directed and habitual behavior through successor representation of states*** A great mystery had been that how model-based and model-free RLs, whose typical algorithms are so different in formulae, can be both hosted by corticostriatal-DA circuits, different parts of which should still share basic architectures. Recent work (Russek *et al.*, 2017; Gershman, 2018) has provided a brilliant potential solution to this by proposing that certain types of goal-directed (model-based) behavior, having sensitivity to changes in outcome values, can be achieved through a particular type of state representation called the successor representation (Dayan, 1993), combined with the ever-suggested representation of RPE by DA (Montague *et al.*, 1996; Schultz *et al.*, 1997). In the successor representation, individual states are represented by a sort of closeness to their successor states, or more accurately, by time-discounted cumulative future occupancies of these states. Behavior based on this representation is not fully goal-directed, having difficulty in revaluation of state transition or policy, which has been demonstrated in actual human behavior (Momennejad *et al.*, 2017) referred to as “subtler, more cognitive notion of habit” by the authors (Momennejad *et al.*, 2017). Successor representation and value update based on it have been suggested to be implemented in the prefrontal/hippocampus-dorsomedial/ventral striatum circuits (Garvert *et al.*, 2017; Russek *et al.*, 2017; Stachenfeld *et al.*, 2017), while circuits including dorsolateral striatum might implement habitual or model-free behavior through “punctate” (i.e., individual) representation of states or actions.
4. ***Sustained DA response to predictable reward, possibly related to state representation*** The original experiments that led to the proposal of representation of RPE by DA (Montague *et al.*, 1996; Schultz *et al.*, 1997) have shown that DA response to reward disappears after monkeys repeatedly experienced the stimulus(-action)-reward association and the reward presumably became predictable for them. However, sustained, and often ramping, dopamine signals to/towards (apparently) predictable reward has been widely observed in recent years (Howe *et al.*, 2013; Collins *et al.*, 2016; Hamid *et al.*, 2016; Hamid *et al.*, 2019; Kim *et al.*, 2019; Mohebi *et al.*, 2019; Guru *et al.*, 2020; Sarno *et al.*, 2020). There are a number of possible accounts for such sustained DA signals, positing that they represent RPE (Gershman, 2014; Morita & Kato, 2014; Kato & Morita, 2016; Kim *et al.*, 2019; Mikhael *et al.*, 2019; Song & Lee, 2020) or something different from RPE (Howe *et al.*, 2013; Hamid *et al.*, 2016; Hamid *et al.*, 2019; Mohebi *et al.*, 2019; Guru *et al.*, 2020; Sarno *et al.*, 2020) or both (Lloyd & Dayan, 2015; Collins *et al.*, 2016). Of particular interest to our present work, one hypothesis (Gershman, 2014) suggests that sustained (ramping) DA signals might represent sustained RPE generated due to imperfect approximation of value function in the system using representation of states by low-dimensional features.

Referring to these different streams of findings and suggestions, we propose a computational model of potential mechanisms underlying the difficulty in resisting undesired habitual behavior to obtain reward.

## Results

### Goal-based reduced successor representation of states under non-resistant policy

We modeled a person’s series of actions to obtain a particular reward, such as alcohol, nicotine, or non-substance such as betting ticket, gaming, or social interaction, by a series of modeled person’s actions on a sequence of states from the start state to the goal state, where the reward is given (Fig. 1A). At each state except for the goal state, the person can take either of two actions, “Go”: proceed to the next state, and “No-Go”: stay at the same state (as considered in our previous work (Kato & Morita, 2016) in a different context). We considered a case that the person has long been regularly taking behavior to obtain the reward without resisting temptation. In the model, it corresponds to that the person has long experienced transitions towards the rewarded goal according to a policy that takes only “Go” at any state, which we refer to as the Non-Resistant policy. We assumed that through such long-standing experiences of behavior according to the Non-Resistant policy, the person has established a particular state representation, where each state is represented by the discounted future occupancy of the final successor state, namely, the rewarded goal state, under that policy. Specifically, we considered a single (i.e., scalar) feature *x* and assumed that the *k*-th state, *S*_*k*_ (*k* = 1, …, *n*; *S*_1_ is the start state and *S*_*n*_ is the goal state), is represented by:

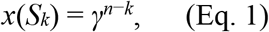

where *γ* is the time discount factor. The number of states (*n*) was set to 10, and the time discount factor (*γ*) was assumed to be 0.97, resulting in that the discounted value at the start state was 0.97^9^ ≈ 0.76 times of the value at the goal, unless otherwise mentioned (Fig. 1B) (see the Materials & Methods for rationale for these parameter values).

**Figure 1.**
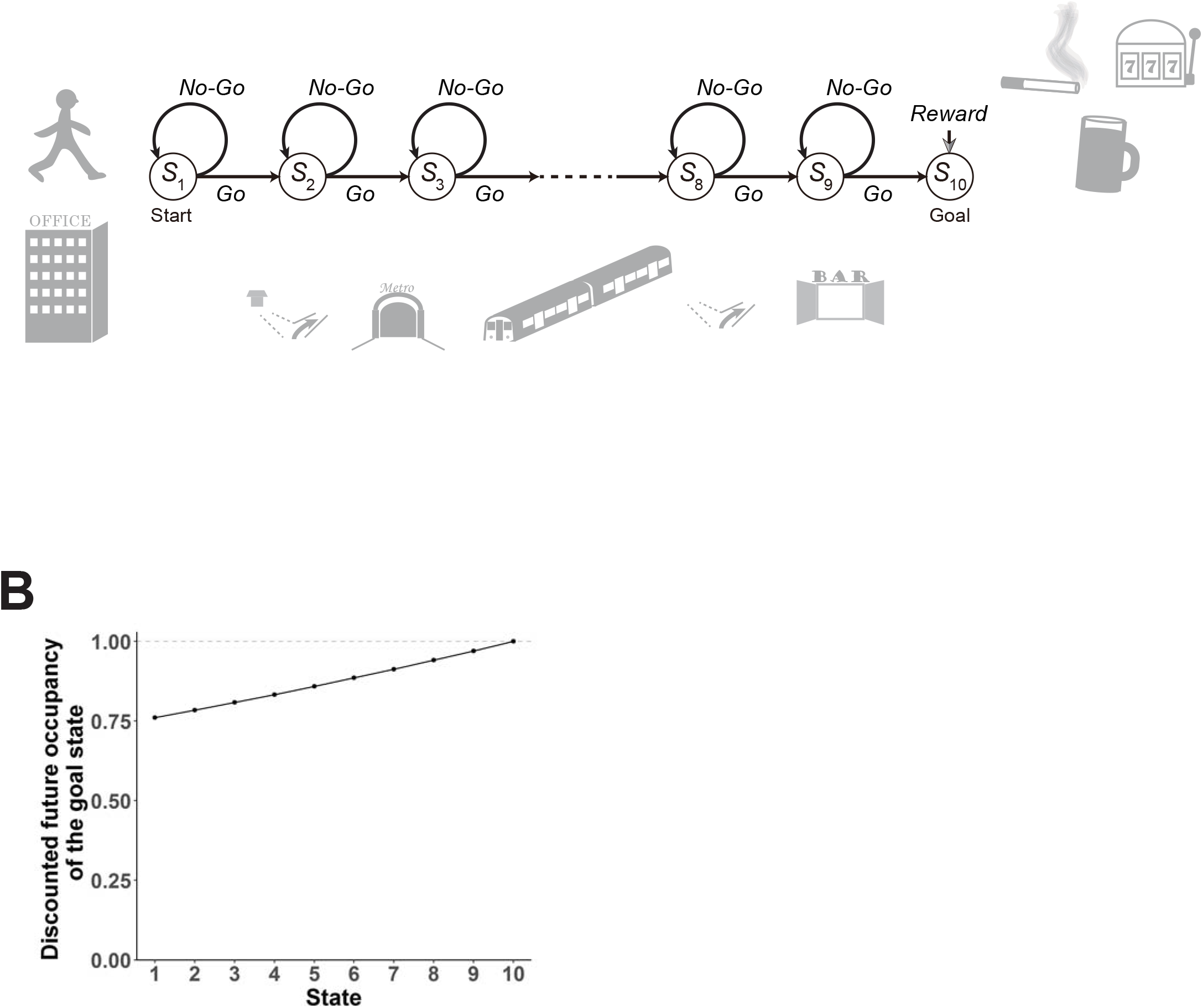
Schematic diagram of the model and the assumed goal-based representation of states under the Non-Resistant policy. **(A)** Schematic diagram of the model, adapted, with alterations, from Fig. 1 of (Kato & Morita, 2016). **(B)** Assumed representation of states by the discounted future occupancy of the final successor state, i.e., the goal state, under the Non-Resistant policy, in which only “Go” is chosen at any state except for the goal state. The vertical axis indicates the discounted future occupancy of the goal state in the case of starting from each state (corresponding to the scalar feature of the state in this representation), given by *x*(*S_k_*) = *γ*^10−*k*^ (Eq. 1) for state *S*_*k*_ (*k* = 1, …, 10; *S*_1_ is the start state and *S*_10_ is the goal state), where *γ* is the time discount factor (set to 0.97 in this figure).

This representation, which we will refer to as the goal-based representation, can be said to be a dimension-reduced version of successor representation; in the genuine successor representation (Dayan, 1993; Russek *et al.*, 2017; Gershman, 2018), every state is represented by a vector of expected cumulative discounted future state occupancies for all the states, whereas in the above goal-based representation, every state is represented by the discounted future occupancy of only the goal state. Because the genuine successor representation requires the number of features equal to the number of states, dimension reduction has been considered (c.f., (Gehring, 2015; Barreto *et al.*, 2016; Gardner *et al.*, 2018)). Given the general suggestion of dimension reduction in state representations in the brain (Gershman & Niv, 2010; Niv, 2019), it would be conceivable that the brain adopts dimension-reduced versions of successor representation, such as the goal-based representation assumed above. Notably, the state value function under the Non-Resistant policy in the case of the assumed structure of state transitions and rewards (Fig. 1A) can be precisely represented as a linear function of the scalar feature of the assumed goal-based representation. Specifically, the state value for *S*_*k*_ is given by

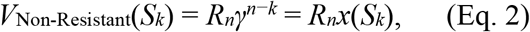

where *R*_*n*_ is the reward value obtained at the goal state, which was assumed to be 1. Therefore, the assumed goal-based representation can be said to be a minimal representation for achieving accurate state values. Moreover, this representation inherits the sensitivity to changes in the reward value at the goal from the genuine successor representation, and thus the agent (person) having acquired this representation remains to be goal-directed in terms of sensitivity to changes in the goal value. It would thus be conceivable that such a goal-based representation has been acquired through long-standing behavior. We will discuss possible implementation of such a representation in the brain, including potentially supporting findings, as well as possible experimental validation of a generalized version of such a representation in the Discussion.

### RPEs under resistant policy, with state representation under non-resistant policy

We then modeled a situation where the person decides to attempt cessation of the habitual reward-obtaining behavior by assuming that the person starts to take a new policy, referred to as the Resistant policy, in which not only “Go” but also “No-Go” action is chosen with a certain probability, *P*_No-Go_, at each state preceding the goal. Crucially, we assumed that the goal-based state representation has been established so rigidly through long-standing behavior under the Non-Resistant policy that the representation does not change after the person changes the policy to the Resistant policy. We therefore assumed that the person tries to approximate the state value function under the new, Resistant policy by a linear function of the abovementioned scalar feature, *x*(*S_k_*), with a coefficient *w*:

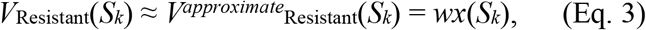

by updating the coefficient *w* using the temporal-difference (TD) RPE at every time step:

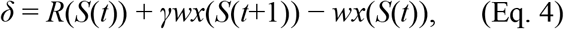

where *S*(*t*) and *S*(*t*+1) are the states at time *t* and *t*+1, respectively, and if *S*(*t*) is the goal state, the term *γwx*(*S*(*t*+1)) is dropped. *R*(*S*(*t*)) is the reward value obtained at *S*(*t*), which was assumed to be 0 except for the goal state. Specifically, *w* was assumed to be updated as follows:

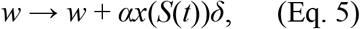

where *α* is the learning rate, which was set to 0.5 unless otherwise mentioned. This way of linear function approximation and TD-RPE-based update (Sutton, 1988; Sutton & Barto, 2018) has been typically assumed in neuro-computational models and is considered to be implementable through synaptic plasticity depending on DA, which represents *δ*, and presynaptic activity, which represents *x*(*S*(*t*)) (Montague *et al.*, 1996; Russek *et al.*, 2017). The initial value of *w* was set to *R*_*n*_ (= 1), with which the approximate value function exactly matches the true value function under the Non-Resistant policy. Notably, we did not simulate the person’s behavior under the Non-Resistant policy, but instead just assumed this initial value of *w* and simulated the person’s behavior under the Resistant policy only. The probability of “No-Go” choice (*P*_No-Go_) was set to 0.75; later we also describe results with different values of *P*_No-Go_.

We then examined RPEs generated upon each decision, “Go” or “No-Go”, at each state before the goal state or upon reaching the goal state. Figure 2Aa shows a single simulation example of RPEs generated at each state in the first episode. In this episode, the person chose “No-Go” once at *S*_3_, twice at *S*_5_ and *S*_6_, four times at *S*_4_, seven times at *S*_1_, *S*_2_, and *S*_9_, nine times at *S*_8_, and never at *S*_7_. The blue crosses indicate RPEs generated upon “Go” decisions, whereas the red crosses indicate the means of RPEs generated upon “No-Go” decisions, and the black cross indicates RPE generated at the goal state. The magenta circles indicate the summation of RPEs generated upon “No-Go” decisions at the same states. As shown in the figure, when the person chose “No-Go”, negative RPEs were generated, whereas theoretically no RPE is generated upon choosing “Go” (though tiny numerical errors existed (the same applies throughout)), and when the person eventually reached the rewarded goal state, a positive RPE was generated. Figure 2Ab shows the mean and standard deviation across simulations. The same features as observed in the example simulation are observed.

**Figure 2.**
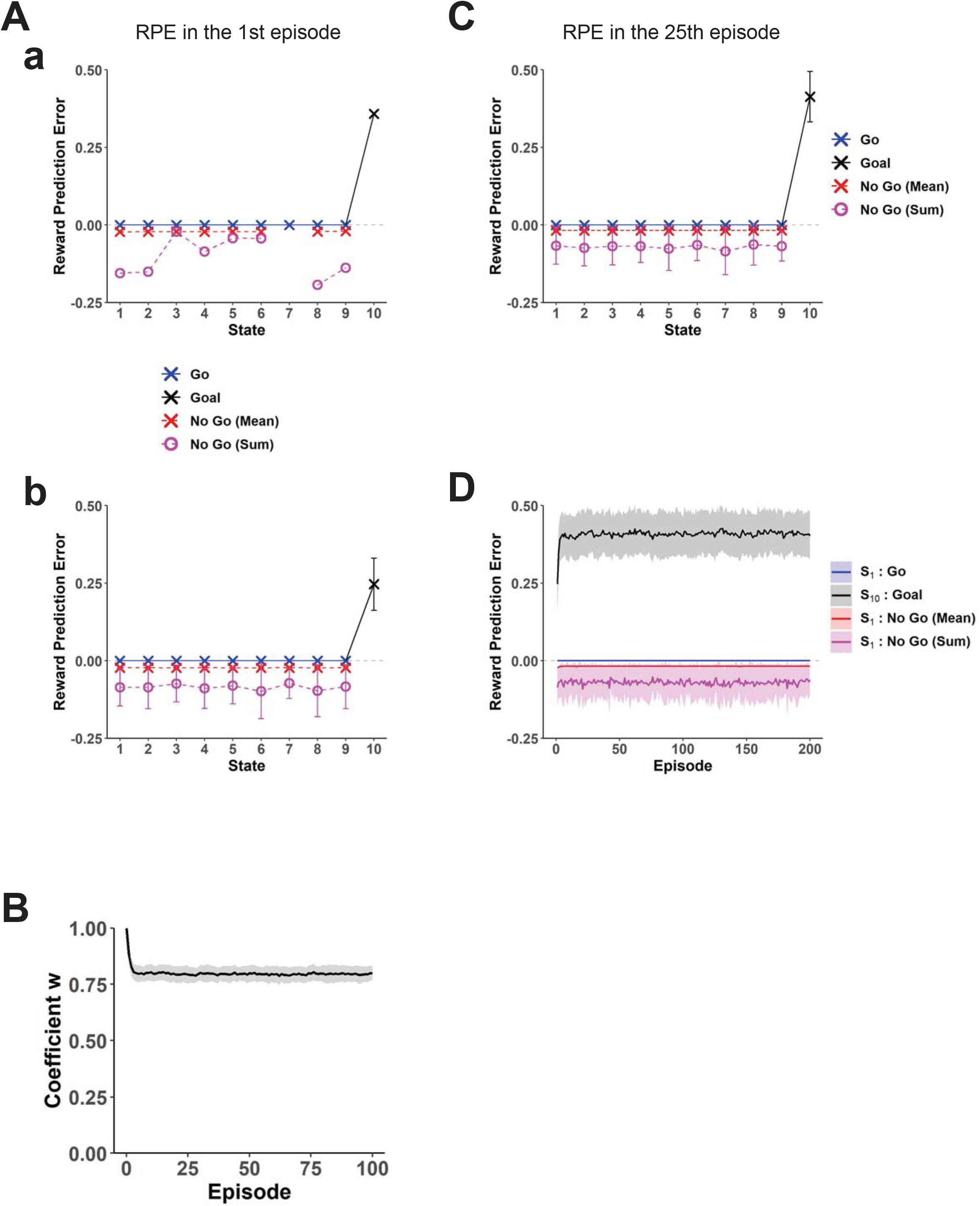
RPEs generated under the Resistant policy, with the goal-based representation under the Non-Resistant policy. **(A) (a)** A single simulation example of RPEs generated at each state in the 1st episode. The blue crosses indicate RPEs generated upon “Go” decisions, whereas the red crosses indicate the means of RPEs generated upon “No-Go” decisions, and the black cross indicates RPE generated at the goal state. The magenta circles indicate the summation of RPEs generated upon “No-Go” decisions at the same states. **(b)** RPEs generated at each state in the 1st episode, averaged across simulations. The error bars indicate ± standard deviation (SD); this is also applied to the following figures. **(B)** Over-episode change of the coefficient *w* of the approximate value function at the end of each episode, averaged across simulations, and the shading indicates ± SD (this is also applied to the following figures); the assumed initial value (*w* = 1) is also plotted at episode = 0 with SD = 0. **(C)** RPEs generated at each state in the 25th episode, where the coefficient *w* has become nearly stationary, averaged across simulations (± SD). **(D)** The changes of RPEs over episodes, averaged across simulations (± SD). RPEs generated upon “Go” decisions (blue) and “No-Go” decisions (mean (red) and summation (magenta) per episode) at the start state, and RPE generated at the goal state (black).

Figure 2B shows the over-episode change of the coefficient *w* of the approximate value function at the end of each episode, averaged across simulations. As shown in the figure, *w* decreases from its initial value, *R*_*n*_ (= 1), and becomes (almost) stationary, meaning that the negative and positive RPE-based updates become overall balanced. We examined RPEs after the coefficient *w* becomes nearly stationary, in particular, in the 25th episode. Figure 2C shows the results averaged across simulations. As shown in the figure, negative RPEs were generated upon “No-Go” decisions, whereas theoretically no RPE is generated upon “Go” decisions, and a large positive RPE was generated upon reaching the rewarded goal state. We also examined how the amplitudes of RPEs change over episodes, specifically for RPEs generated upon “Go” or “No-Go” decisions at the start state and RPE generated at the goal state (Fig. 2D). As shown in the figure, after a few initial episodes, the amplitudes, averaged across simulations, become nearly stationary.

We also examined the cases with different parameters, in particular, with the probability of “No-Go” choice (*P*_No-Go_) varied over 0.5, 0.75 (assumed above), and 0.9, and the time discount factor (*γ*) varied over 0.95, 0.97 (assumed above), and 0.99. Figure 3 shows the results for RPEs in the 25th episode (Fig. 3A) and over-episode changes of RPEs upon “Go” or “No-Go” decisions at the start state and RPE at the goal state (Fig. 3B) (the center panels of Fig. 3A,B show the same data as shown in Fig. 2C,D, respectively). As shown in the figures, basic features mentioned above are largely preserved over these parameter ranges. We further examined the case where the learning rate (*α*), which has so far been set to 0.5, was smaller, specifically *α* = 0.1. Figure 4 shows the results. As shown in the figure, it took more episodes for the coefficient *w* and RPEs to become nearly stationary, as expected, but thereafter the generated RPEs look nearly comparable to the case with the original larger learning rate (Fig. 2C). This is considered to be because the RPEs primarily come from the sustained mismatch between the true state value function and the estimated value function (i.e., linear function of the features), and the estimated value function on average should not so much depend on the learning rate once learning (probabilistically) converges although the amplitude of within-episode changes of *w* depends on the learning rate. As shown in these simulation results with parameters varied, the pattern of generated sustained RPEs appears to be robust to a good extent. These RPEs, in particular the large positive RPE upon goal reaching, are considered to potentially contribute to the difficulty in resisting habitual behavior to obtain reward.

**Figure 3.**
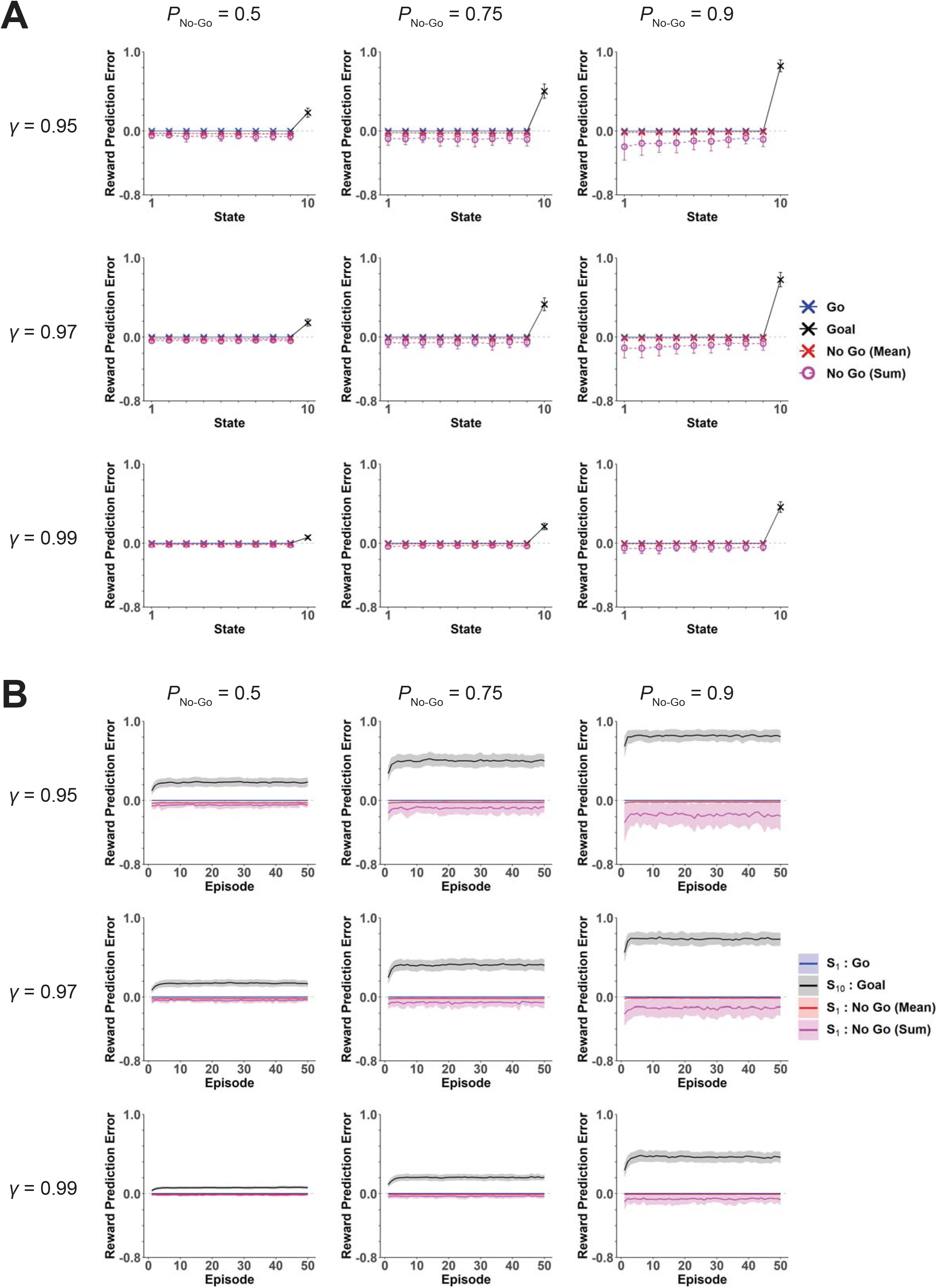
Cases with different parameters. The probability of “No-Go” choice (*P*_No-Go_) was set to 0.5, 0.75 (the value assumed in Fig. 2), and 0.9. The time discount factor (*γ*) was set to 0.95, 0.97 (the value assumed in Fig. 2), and 0.99. **(A)** RPEs generated at each state in the 25th episode, averaged across simulations (± SD). Notations are the same as those in Fig. 2C (blue: “Go”, red cross: “No-Go” mean, magenta circle: “No-Go” sum, black: goal), and the center panel shows the same data as shown in Fig. 2C. **(B)** Over-episode changes of RPEs upon “Go” and “No-Go” decisions at the start state, and RPE generated at the goal state, averaged across simulations (± SD). Notations are the same as those in Fig. 2D (blue: “Go” at the start, red: “No-Go” mean at the start, magenta: “No-Go” sum at the start, black: at the goal), and the center panel shows the same data as shown in Fig. 2D.

**Figure 4.**
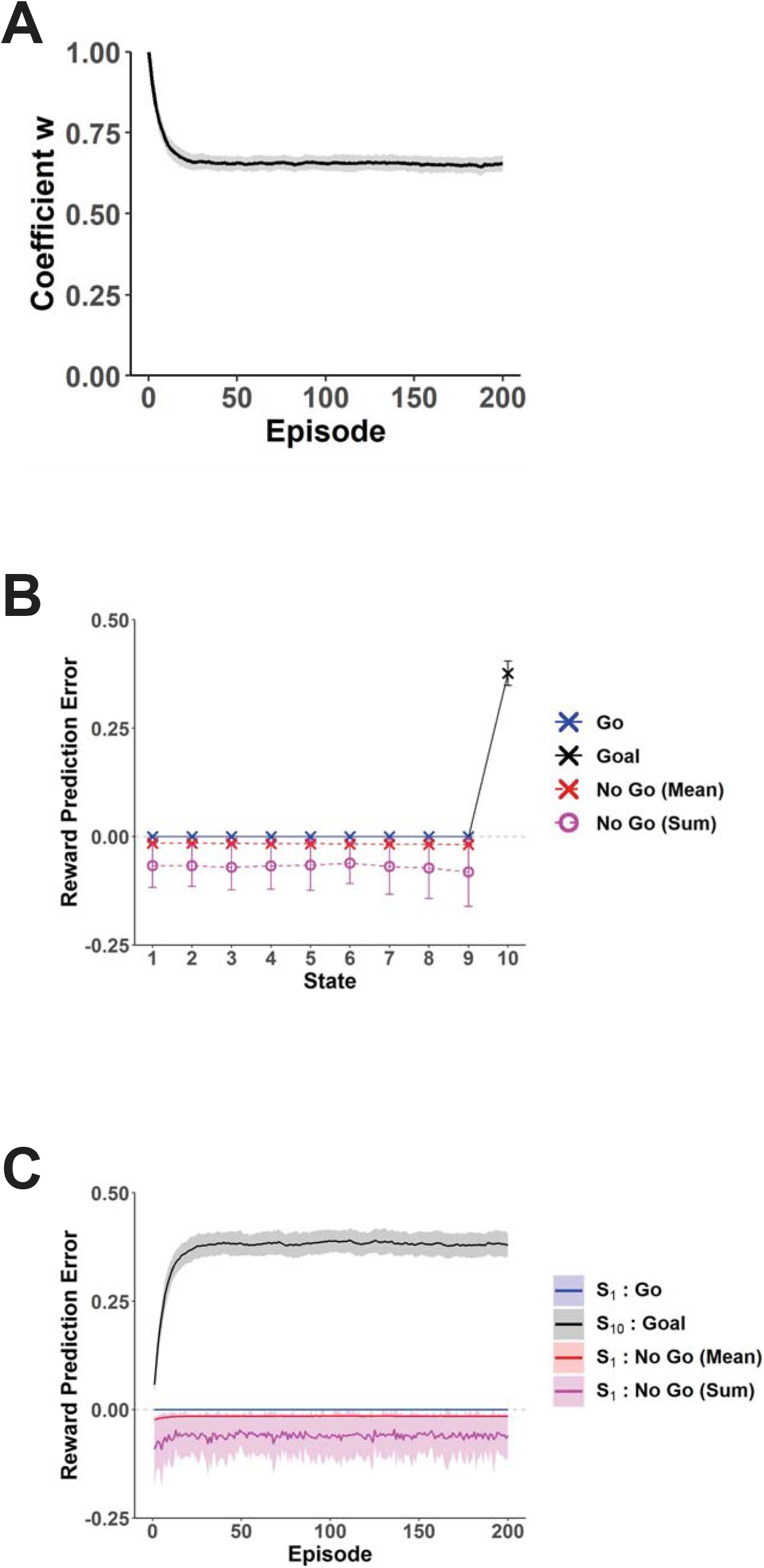
Case where the learning rate was smaller (*α* = 0.1) than the original cases (*α* = 0.5). The probability of “No-Go” choice (*P*_No-Go_) and the time discount factor (*γ*) were set to 0.75 and 0.97, respectively (the values assumed in Fig. 2). **(A)** Over-episode change of the coefficient *w* of the approximate value function at the end of each episode, averaged across simulations, and the shading indicates ± SD. Notations are the same as those in Fig. 2B. **(B)** RPEs generated at each state in the 25th episode, averaged across simulations (± SD). Notations are the same as those in Fig. 2C and Fig. 3A. **(C)** Over-episode changes of RPEs upon “Go” and “No-Go” decisions at the start state, and RPE generated at the goal state, averaged across simulations (± SD). Notations are the same as those in Fig. 2D and Fig. 3B.

If the person continues to take the Resistant policy for a number of times, it can be expected that the goal-based representation of states established under the Non-Resistant policy slowly changes, and gradually approaches the representation under the Resistant policy, particularly through TD learning of state representation itself (Gershman *et al.*, 2012; Gardner *et al.*, 2018). We thus examined how RPEs become if it occurs. Specifically, we again assumed the original parameters considered in Fig. 2, i.e., *P*_No-Go_ = 0.75, *γ* = 0.97, and *α* = 0.5, and conducted simulations in the same way as above, except that this time we also updated the scalar feature of the state (i.e., *x*(*S*(*t*))) at every time step by using the TD error of the goal-based representation:

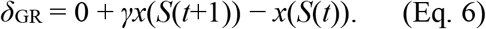

Specifically, the scalar feature was updated as follows:

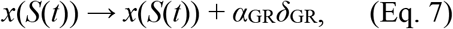

where *α*_GR_ is the learning rate for this update and was set to 0.05, except for the goal state, for which the TD error of the goal-based representation should be theoretically 0 and thus no update was implemented. Figure 5A shows the scalar feature of each state (i.e., *x*(*S_k_*)) after 50, 100, and 200 episodes (black dotted, dashed, and solid lines, respectively), averaged across simulations, in comparison to the original ones (gray line). As shown in the figure, the curve became steeper as episodes proceeded. This is considered to reflect that longer time is required, on average, for goal reaching under the Resistant policy than under the Non-Resistant policy and thus the expected discounted future occupancy of the goal state should be smaller for the Resistant policy. Figure 5B shows the RPEs generated in the 200th episode, averaged across simulations, and Figure 5C shows the over-episode changes in the RPEs generated upon “Go” or “No-Go” decisions at the start state and the RPE generated at the goal state. As shown in these figures, the large positive RPE generated upon goal reaching observed in the case with the original state representation (Fig. 2C) gradually decreased, while positive RPEs with smaller amplitudes gradually appeared upon “Go” decisions in the states other than the goal. This indicates that if the large positive RPE upon goal reaching is especially harmful for cessation of habitual reward-seeking, it could be resolved if the person does not give up resisting temptation, even with the help of clinical intervention such as alternative reward upon “No-Go” choices, until the state representation considerably changes.

**Figure 5.**
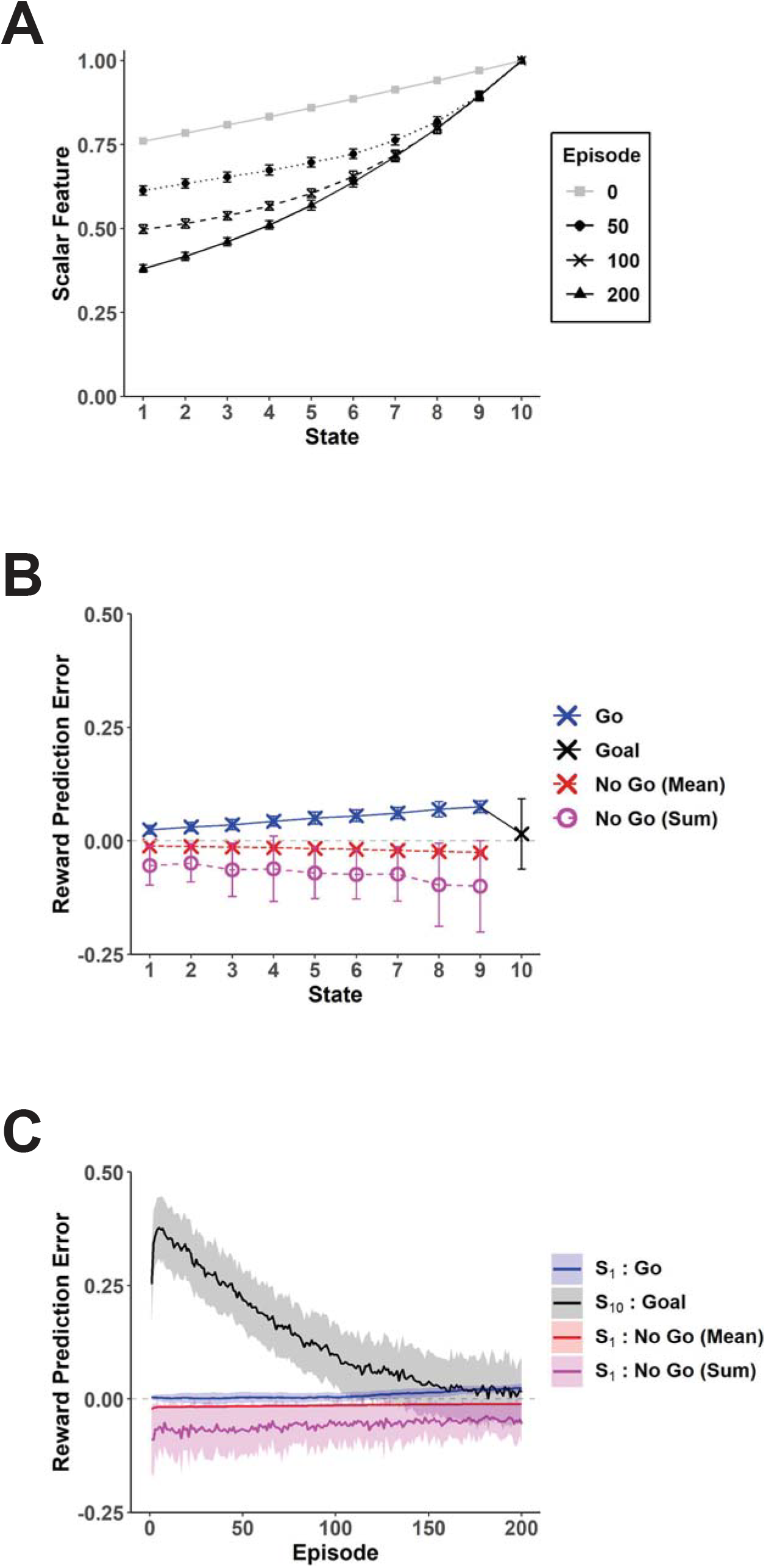
RPEs generated under the Resistant policy, in the case where the goal-based state representation itself slowly changed and approached the goal-based representation under the Resistant policy. The probability of “No-Go” choice (*P*_No-Go_) was set to 0.75, and the time discount factor (*γ*) was set to 0.97. The learning rate for the update of the coefficient *w* of the approximate value function (*α*) was set to 0.5, and the learning rate for the update of the scalar feature of states in the goal-based representation (*α*_GR_) was set to 0.05. **(A)** Scalar feature of each state (i.e., *x*(*S_k_*)) after 50, 100, and 200 episodes (black dotted, dashed, and solid lines, respectively), averaged across simulations (± SD), in comparison to the original ones (gray line) that are the same as those shown in Fig. 1B. **(B)** RPEs generated at each state in the 200th episode, averaged across simulations (± SD). Notations are the same as those in Fig. 2C (blue: “Go”, red cross: “No-Go” mean, magenta circle: “No-Go” sum, black: goal). **(C)** Over-episode changes of RPEs upon “Go” and “No-Go” decisions at the start state, and RPE generated at the goal state, averaged across simulations (± SD). Notations are the same as those in Fig. 2D (blue: “Go” at the start, red: “No-Go” mean at the start, magenta: “No-Go” sum at the start, black: at the goal).

### Comparison to the cases with punctate representation or genuine successor representation of states

For comparison, we considered a case where each state is represented in the “punctate” manner (i.e., individually). For this case, we assumed that each state has its own state value, *V*_punctate_(*S_k_*), and it is updated using TD-RPE in the punctate system, *δ*_punctate_:

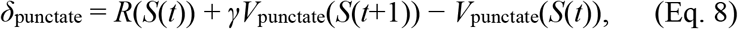

where if *S*(*t*) is the goal state, the term *γV*_punctate_(*S*(*t*+1)) is dropped. Specifically, *V*_punctate_ was assumed to be updated as follows:

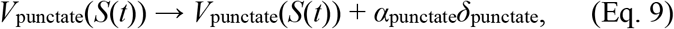

where *α*_punctate_ is the learning rate for the punctate system, which was set to 0.5. The initial values for the punctate state values were assumed to be:

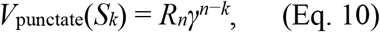

which are the state values under the Non-Resistant policy as considered in Eq. 2.

Figure 6 shows the RPEs generated in the punctate system in the same various conditions as examined for the system with the goal-based representation (Fig. 3). Comparing these two figures, prominent differences are that whereas theoretically no RPE occurs upon “Go” decisions and a large positive RPE is generated upon goal reaching in the case with the goal-based representation, (relatively small) positive RPEs are generated upon “Go” decision and theoretically no RPE occurs at the goal in the case with punctate representation. These differences are considered to reflect different characteristics of updates done with the different ways of state representation. Specifically, in the case with the goal-based representation, only the coefficient of approximate value function was updated and the state representation established under the Non-Resistant policy was (assumed to be) unchanged, resulting in sustained mismatch between the true and approximate value functions. In contrast, in the case with punctate state representation, the value of each state was directly updated so that there is no such sustained mismatch.

**Figure 6.**
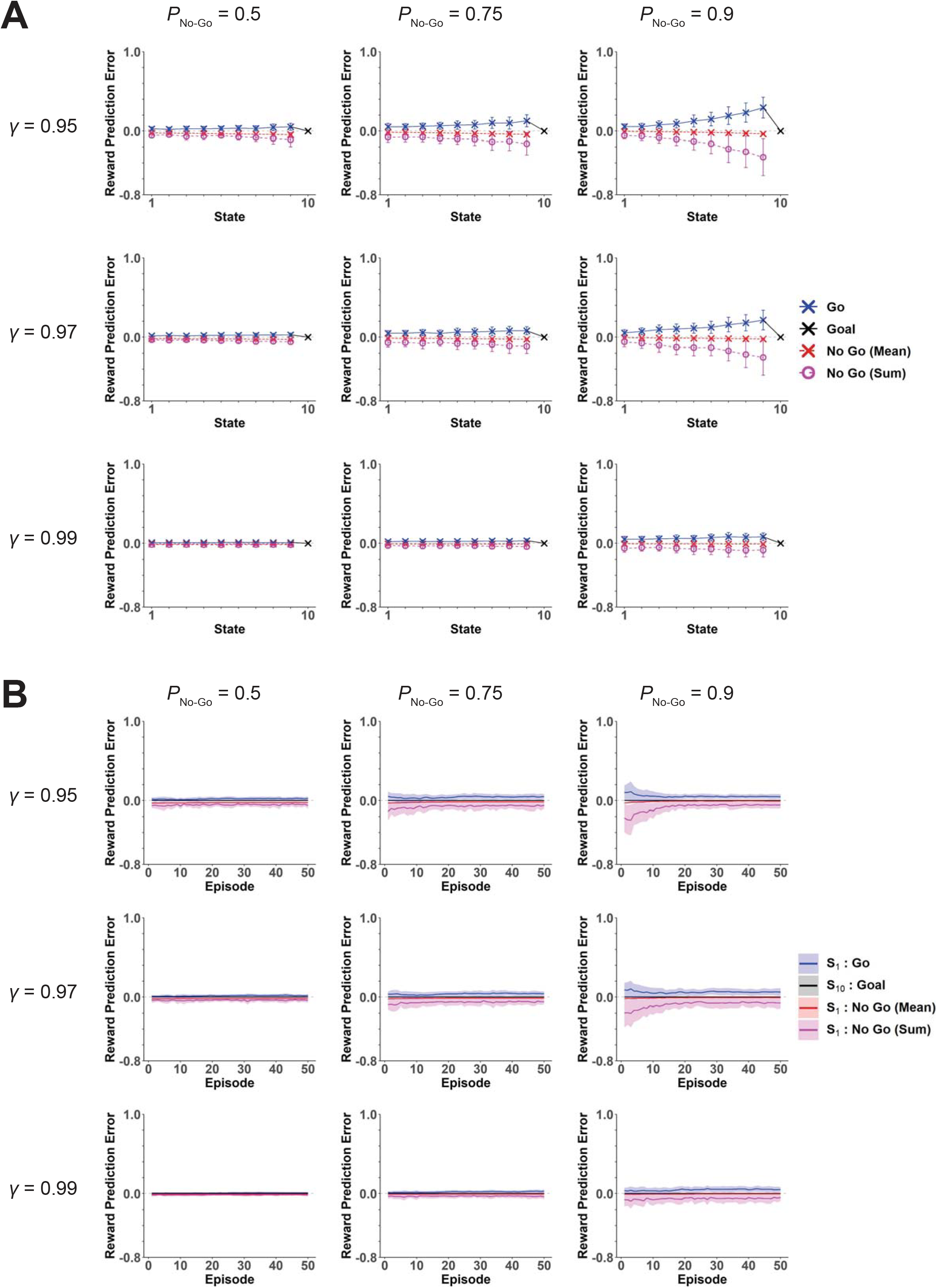
RPEs generated in the system with punctate representation of states. The conditions, parameters, and notations are the same as those in Fig. 3. **(A)** RPEs generated at each state in the 25th episode, averaged across simulations (± SD) (blue: “Go”, red cross: “No-Go” mean, magenta circle: “No-Go” sum, black: goal). **(B)** Over-episode changes of RPEs upon “Go” and “No-Go” decisions at the start state, and RPE generated at the goal state, averaged across simulations (± SD) (blue: “Go” at the start, red: “No-Go” mean at the start, magenta: “No-Go” sum at the start, black: at the goal).

We also considered a case where the states are represented by the genuine successor representation. Specifically, we assumed that each state *S*_*k*_ is represented by *n* features *x_j_*(*S_k_*) (*j* = 1, …, *n*) indicating the time-discounted future occupancy of *S*_*j*_ under the Non-Resistant policy:

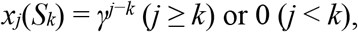

and the value function under the Resistant policy is approximated by a linear function of them:

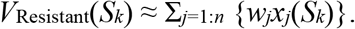

The coefficients *w*_*j*_ (*j* = 1, …, *n*) are updated by using the TD RPE:

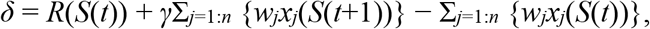

where the middle term including *S*(*t*+1) is dropped if *S*(*t*) is the goal state, according to the following rule:

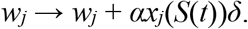

The initial values of *w*_*j*_ were set to 0 for *j* = 1, …, *n*−1 and *R*_*n*_ (= 1) for *j* = *n*, with which the approximate value function exactly matches the true value function under the Non-Resistant policy. Figure 7A shows the RPEs generated at each state in the 25th episode, and Figure 7B shows the over-episode changes in the RPEs generated upon “Go” or “No-Go” decisions at the start state and the RPE generated at the goal state, both with the original parameters considered in Fig. 2, i.e., *P*_No-Go_ = 0.75, *γ* = 0.97, and *α* = 0.5. As shown in the figures, the patterns of RPEs are similar to those in the case of punctate representation (the center panels of Fig. 6A,B) and differ from those in the case of the goal-based representation. Figure 7C shows the coefficients *w*_*j*_ of the approximate value function after the 1st episode (Fig. 7Ca) and 25th episode (Fig. 7Cb), and Figure 7D shows the over-episode changes of the coefficients for the features corresponding to the start state (red line), the state preceding the goal (*S*_9_) (blue line), and the goal state (black line). As shown in these figures, the coefficients for the features corresponding to the states preceding the goal became negative. It is considered that because of these negative coefficients, the true value function under the Resistant policy could be well approximated even by a linear function of the features (discounted occupancies) under the Non-Resistant policy.

**Figure 7.**
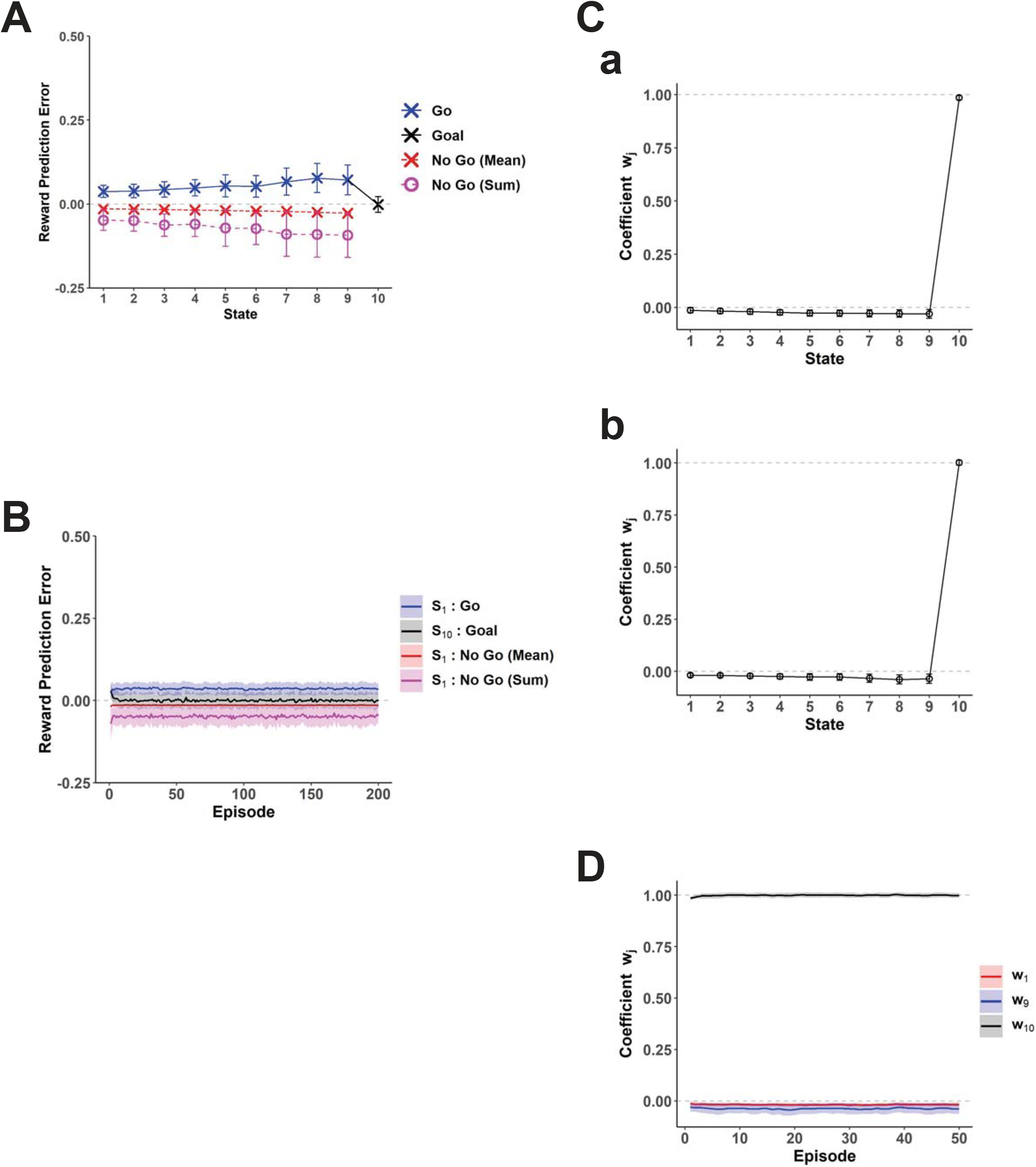
Case with the genuine successor representation. The probability of “No-Go” choice (*P*_No-Go_) was set to 0.75, and the time discount factor (*γ*) was set to 0.97. **(A)** RPEs generated at each state in the 25th episode, averaged across simulations (± SD). Notations are the same as those in Fig. 2C (blue: “Go”, red cross: “No-Go” mean, magenta circle: “No-Go” sum, black: goal). **(B)** Over-episode changes of RPEs upon “Go” and “No-Go” decisions at the start state, and RPE generated at the goal state, averaged across simulations (± SD). Notations are the same as those in Fig. 2D (blue: “Go” at the start, red: “No-Go” mean at the start, magenta: “No-Go” sum at the start, black: at the goal). **(C)** Coefficients *w*_*j*_ of the approximate value function after the 1st episode (a) and 25th episode (b), averaged across simulations (± SD). **(D)** Over-episode changes of the coefficients *w*_*j*_ for the features corresponding to the start state (red line), the state preceding the goal (*S*_9_) (blue line), and the goal state (black line), averaged across simulations (± SD).

### Influence of the goal-based representation system on the punctate action representation system

So far we have examined the cases with different ways of state representation separately. However, as mentioned in the Introduction, it is suggested that there exist multiple value learning systems in the brain, with the system employing successor representation residing in the prefrontal/hippocampus-dorsomedial/ventral striatum circuits, whereas another system adopting punctate representation might locate in the circuits including dorsolateral striatum. Moreover, there are anatomical suggestions of ventral-to-dorsal spiral influences in the striatum-midbrain system (Haber *et al.*, 2000; Joel & Weiner, 2000), and theoretical proposals that such a spiral circuit implements heterarchical reinforcement learning (Haruno & Kawato, 2006) and that the bias of RPE due to drug-induced DA accumulates through the spiral circuit and causes undesired compulsive drug taking in long-term addicts (Keramati & Gutkin, 2013). Inspired by these, we also examined a case with multiple representation/learning systems. Specifically, we assumed that the prefrontal/hippocampus-dorsomedial/ventral striatum circuits adopt the goal-based reduced successor representation of states (rather than the genuine successor representation) whereas the circuits including dorsolateral striatum adopt a punctate representation of individual actions, i.e., “Go” and “No-Go”. This latter assumption was made based on the suggestions that the dorsal/dorsolateral striatum is involved in value learning with actions (O’Doherty *et al.*, 2004; Takahashi *et al.*, 2008). We then assumed that the information of the RPEs generated in the system with the goal-based state representation flows into the system with punctate (i.e., individual) action representation through the spiral circuit (Fig. 8A). Critically, different from the abovementioned previous model (Keramati & Gutkin, 2013), which assumed that the value of the upcoming state but not of the previous state in the ventral circuit flows into the dorsal circuit, we assumed that the information of the values of both upcoming and previous states used for TD RPE calculation originates from the striatum, and effectively the entire TD RPE in the ventral circuit flows into the dorsal circuit (in this regard, somewhat similar assumption was made in (Takahashi *et al.*, 2008)). If both the upcoming and previous values are sent via the direct striatum-midbrain connections, e.g. through the matrix and patch/striosomal neurons as referred to in (Morita *et al.*, 2012), the suggested spiral connections could also convey both information, though it needs to be verified. If either value (or both) is sent to the midbrain via the indirect pathway through the globus pallidus or ventral pallidum, as proposed in (Houk *et al.*, 1995; Doya, 2000; Morita *et al.*, 2012; Morita & Kawaguchi, 2019), our assumption requires spiral connectivity for both direct and indirect pathways.

**Figure 8.**
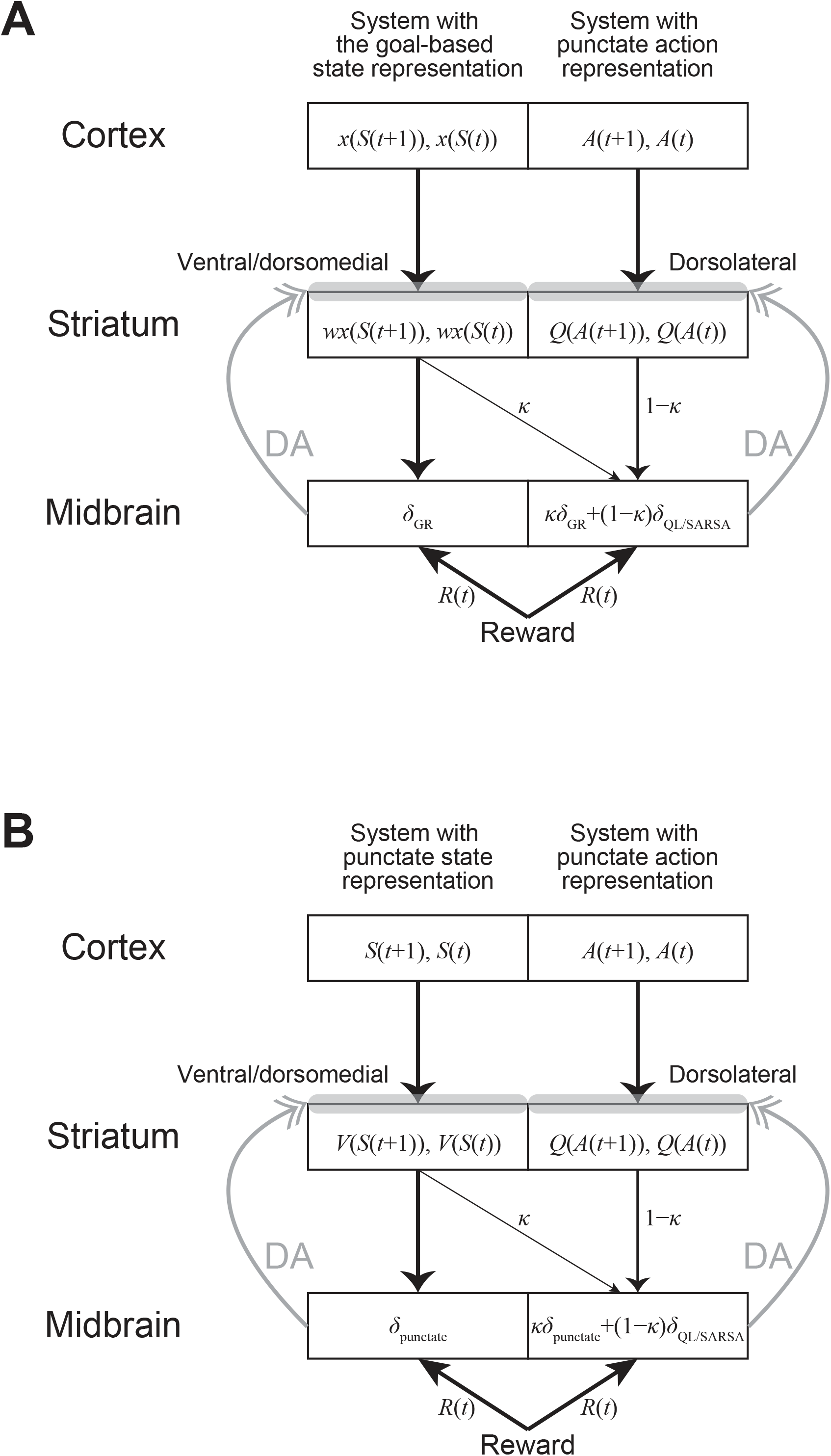
Schematic illustration of the spiral striatum-midbrain circuit and the hypothesized influence of the RPE generated in the circuit including the ventral/dorsomedial striatum on the circuit including the dorsolateral striatum. **(A)** The case where the ventral/dorsomedial circuit employs the goal-based state representation whereas the dorsolateral circuit adopts the punctate (i.e., individual) action representation. In the ventral/dorsomedial circuit (left), the cortex represents the features (discounted future occupancies of the goal state) of the upcoming and previous states and the striatum represents their approximate state values obtained by the linear function of the features with the coefficient *w*. In the dorsolateral circuit (right), the cortex represents the upcoming and previous actions and the striatum represents their action values. The oblique line indicates the spiral striatum-midbrain projections, and *κ* (≥ 0) is a parameter representing the degree of the effect of the RPE generated in the ventral/dorsomedial circuit on the dorsolateral circuit. **(B)** The case where the ventral/dorsomedial circuit employs the punctate (i.e., individual) state representation while the dorsolateral circuit adopts the punctate action representation. In the ventral/dorsomedial circuit (left), the cortex represents the upcoming and previous states and the striatum represents their state values.

We also noticed that a recent study specifically suggested a function of the hierarchical cortico-basal ganglia circuits in habit learning (Baladron & Hamker, 2020), but our model considers a different mechanism.

Regarding the punctate action representation system, we assumed that “Go” and “No-Go” at each state other than the goal state are represented in a punctate manner (i.e., individually) and their values (i.e., action values) are updated by using a combination of the TD RPEs generated in the goal-based state representation system and the TD RPEs of action values. As for the latter, we considered two types: the Q-learning-type and the SARSA-type, both of which have been suggested to be represented by DA (Morris *et al.*, 2006; Roesch *et al.*, 2007). Specifically, we considered the action values

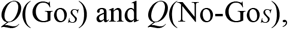

for “Go” and “No-Go” at state *S* (= *S*_1_, …, *S_n_*_−1_), respectively, and considered the TD RPE of Q-learning type:

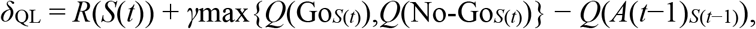

where “max” is the operation to take the maximum and *A*(*t*−1)_*S*(*t*−1)_ is the action actually taken at state *S*(*t*−1), or the TD RPE of SARSA-type:

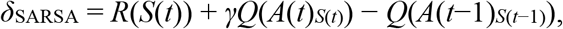

where *A*(*t*)_*S*(*t*)_ is the action actually chosen at state *S*(*t*). For both types, if *S*(*t*) is the goal state, the middle term is dropped, and if *t* is the initial time step within an episode, the last term is dropped. The value of the previous action, *Q*(*A*(*t*−1)_*S*(*t*−1)_), was then assumed to be updated by a combination of either of these RPEs and the RPE generated in the system with the goal-based reduced successor representation of states:

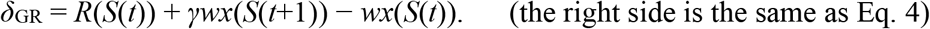

In particular, *Q*(*A*(*t*−1)_*S*(*t*−1)_) was assumed to be updated as:

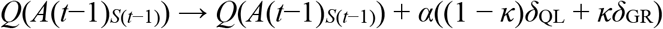

or

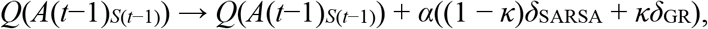

where *κ* (≥ 0) is a parameter representing the degree of the effect of the RPE generated in the system with the goal-based representation, which was varied to be 0, 0.2, or 0.4. We assumed that these TD RPE calculations and updates are implemented in the circuits shown in Fig. 8A. Notably, as appeared in the above equations, we assumed that the TD RPEs containing the reward at *S*(*t*) are used to update the value of action taken at *t*−1 (rather than at *t*). Initial values of the action values for “Go” were set to be the theoretical true values under the Non-Resistant policy, specifically,

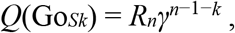

and initial values of the action values for “No-Go” were set to be the values that were one time-step discounted from the initial values for “Go” at the same states:

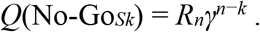

The parameters for the probability of “No-Go” choice, temporal discounting, and learning rate were set to be the original values considered in Fig. 2, i.e., *P*_No-Go_ = 0.75, *γ* = 0.97, and *α* = 0.5.

Figure 9Aa shows examples of across-episode changes of the values of “Go” and “No-Go” at the start state, the middle (5-th) state, and the pre-goal (9-th) state in the case with *κ* = 0 (left panels), 0.2 (middle panels), and 0.4 (right panels) in single simulations with the Q-learning-type TD RPE. When there was no effect of the RPE generated in the system with the goal-based representation (*κ* = 0), all the action values look unchanged from their initial values, as theoretically expected. As the effect of the RPE from the goal-based representation system increased (*κ* = 0.2 and *κ* = 0.4), the values of “Go” and “No-Go” at the start and middle states initially decreased, but the values of “Go” and “No-Go” at the pre-goal state increased, and eventually the action values at all these three states became larger than the values in the case without the effect of the RPE from the goal-based representation system. The initial decreases of action values at the start and middle states are considered to be because of the negative RPEs upon “No-Go” choices in the goal-based representation system, whereas the eventual increases of action values are considered to come from the large positive RPE upon goal-reaching in the goal-based representation system.

**Figure 9.**
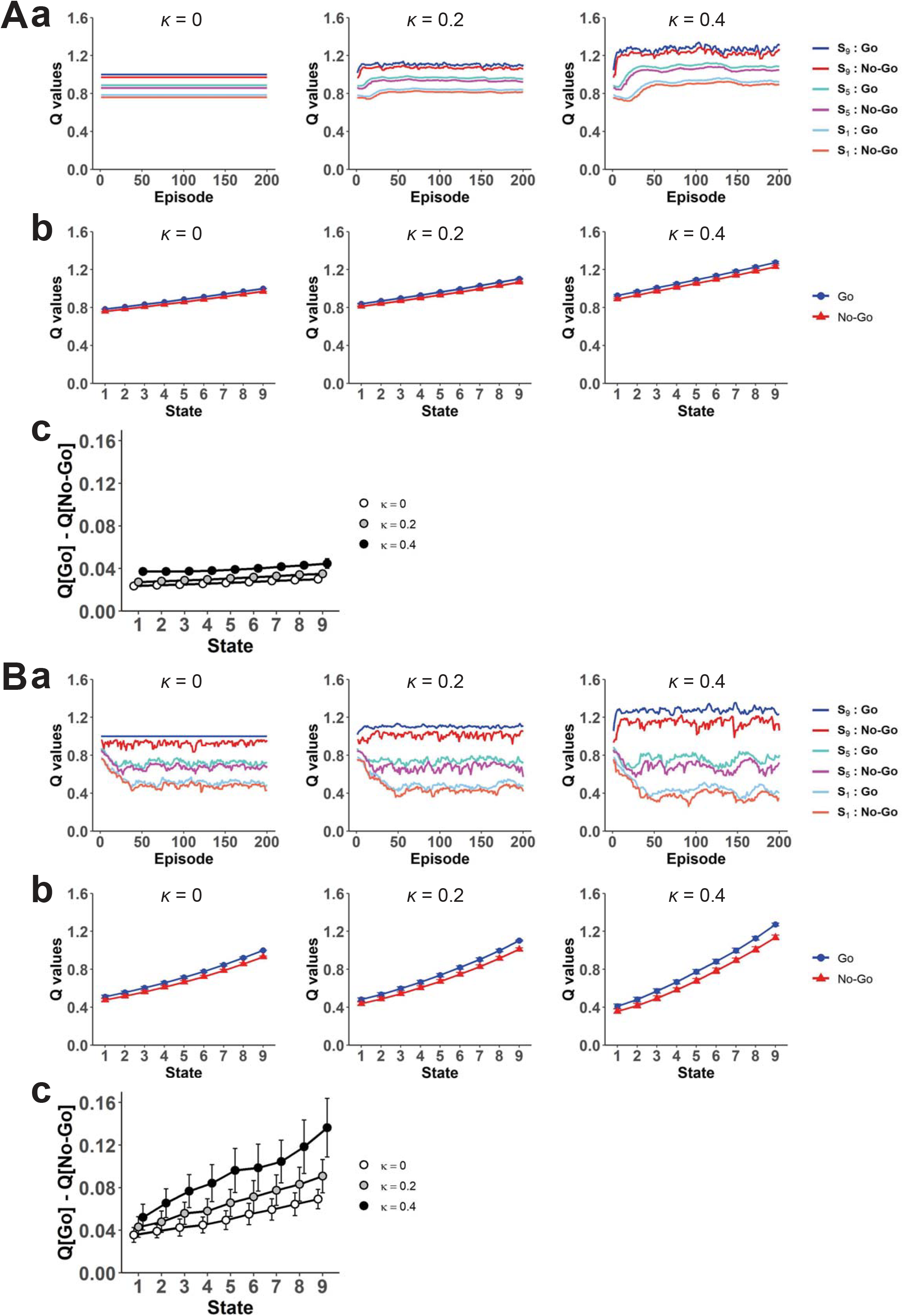
Influence of the RPE generated in the system with the goal-based state representation to the system with punctate action representation. **(A)** Results with the Q-learning-type TD RPE of action values. **(a)** Examples of across-episode changes of the values of “Go” and “No-Go” at the start state, the middle (5-th) state, and the pre-goal (9-th) state in the case with different degrees of the effect of the TD RPE generated in the system with the goal-based state representation (*κ* = 0 (left panels), 0.2 (middle panels), and 0.4 (right panels)) in single simulations. **(b)** The values of “Go” (blue lines) and “No-Go” (red lines) at each state in the case with *κ* = 0 (left panels), 0.2 (middle panels), and 0.4 (right panels), averaged across the 41st to 60th episodes and also across simulations. The error bars indicate ± SD across simulations. **(c)** The differences of the values of “Go” and “No-Go” at each state (*Q*(Go_*Sk*_) − *Q*(No-Go_*Sk*_)) in the case with *κ* = 0, 0.2, and 0.4, averaged across the 41st to 60th episodes and also across simulations. The error bars indicate ± SD across simulations **(B)** Results with the SARSA-type TD RPE of action values. Configurations are the same as those in (A).

Figure 9Ba shows examples of the “Go” and “No-Go” values in single simulations with the SARSA-type TD RPE. When there was no effect of the RPE generated in the system with the goal-based representation (*κ* = 0), the values of actions except for “Go” at the pre-goal state generally decreased from their initial values. This is reasonable, because the on-policy values of these actions under the Resistant policy should be smaller than the values under the Non-Resistant policy due to extra time steps required for goal reaching. As the effect of the RPE from the goal-based representation system increased (*κ* = 0.2 and *κ* = 0.4), the values of “Go” and “No-Go” at the pre-goal state increased, presumably due to the large positive RPE upon goal reaching in the goal-based representation system, while the effects on the action values at the start and middle states appear to be more mixed.

Figure 9Ab shows the values of “Go” and “No-Go” at each state, and Figure 9Ac shows their differences (*Q*(Go_*Sk*_) − *Q*(No-Go_*Sk*_)), in the case with *κ* = 0, 0.2, and 0.4, averaged across the 41st to 60th episodes and also across simulations, with the Q-learning-type TD RPE. Figure 9Bb and 9Bc show the results with the SARSA-type TD RPE. As shown in these figures, in both cases with the different types of TD RPE of action values, the values of “Go” were on average larger than the values of “No-Go”, and the value difference on average increased as the effect of the RPE from the goal-based representation system increased (*κ* = 0.2 and *κ* = 0.4), although there were large variations in the case of the SARSA-type TD RPE. Therefore, if the “Go” and “No-Go” values were assumed to affect the agent’s choice propensity, which was in reality predetermined to be a fixed probability (*P*_No-Go_ = 0.75) in our model as described above, the RPE information flowing from the goal-based representation system to the punctate action representation system through the spiral circuit could potentially enhance deterioration of the resistance to temptation. This result is intuitively understandable because, from the standpoint of the punctate action representation system, the incoming positive RPE from the goal-based representation system upon goal reaching would act as an extra reward.

For comparison, we also examined the case where the information of the RPEs generated in the system with the punctate *state* representation (considered in the previous section), rather than the system with the goal-based state representation, flows into the system with punctate *action* representation (Fig. 8B). Specifically, we conducted simulations of a model that was the same as the one described above except that *δ*_GR_ (RPE in the goal-based representation system) was replaced with *δ*_punctate_ (RPE in the punctate state representation system, Eq. 8 in the previous section). Figure 10 shows the results. Different from the case with the RPE influence from the goal-based representation system, the RPE influence from the punctate state representation system did not increase but rather decreased the differences between the “Go” and “No-Go” values, in both cases with Q-leaning type (Fig. 10Ac) or SARSA-type (Fig. 10Bc) TD RPE of action values. As shown before (Fig. 6), in the system with punctate state representation, negative and positive RPEs are generated upon “No-Go” and “Go” choices, respectively. However, through the influence to the punctate action representation system, these RPEs are used for updating the value of the *previous* action, which is either “No-Go” at the same state or “Go” at the preceding state (except when the agent is at the start state). Therefore, the negative and positive RPEs upon “No-Go” and “Go” choices in the punctate state representation system do not directly affect the values of chosen actions themselves. The simulation results (Fig. 10) indicate that the spiraling RPE influence from the goal-based representation system, but not from the punctate state representation system, could potentially deteriorate the resistance to temptation.

**Figure 10.**
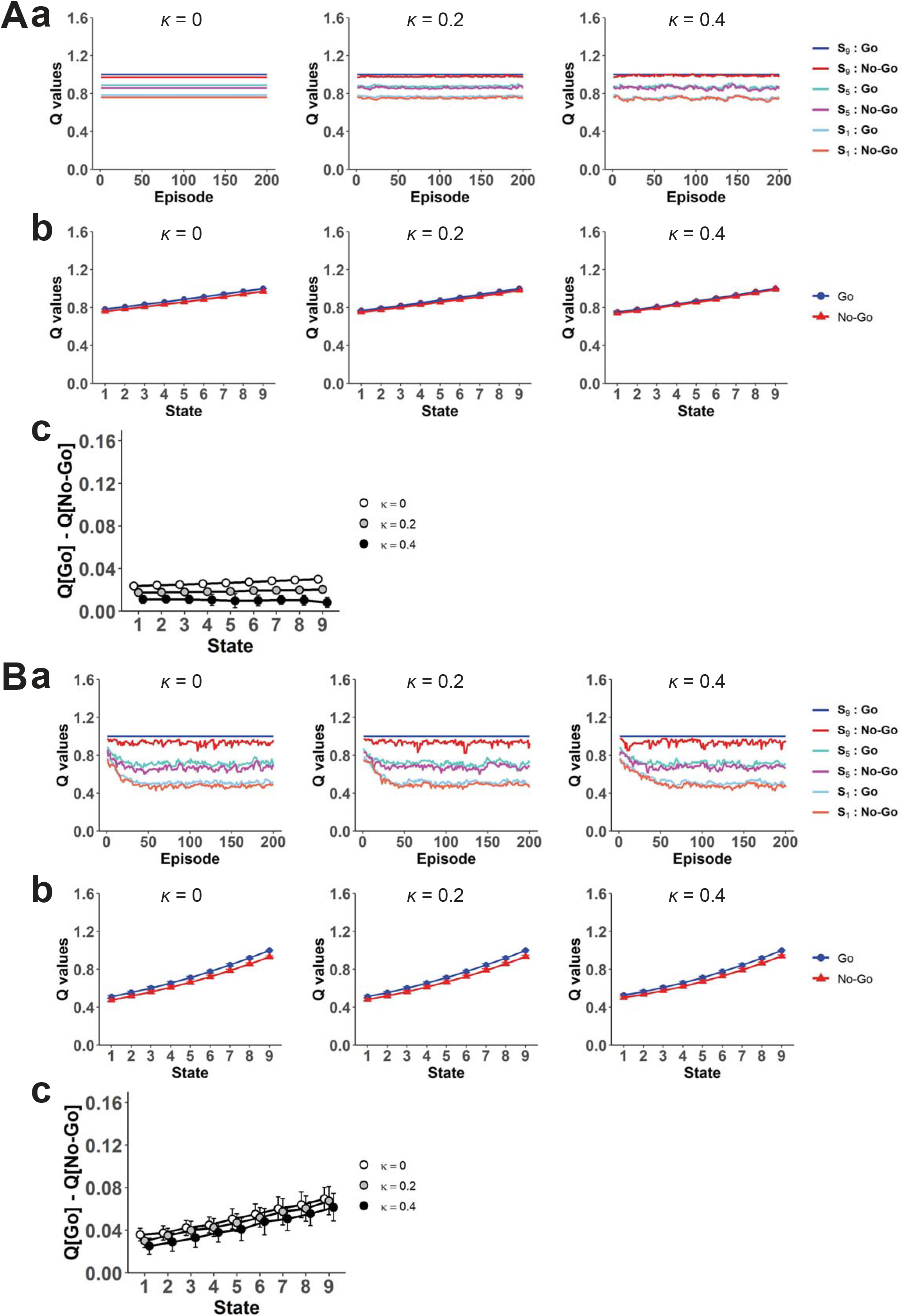
Influence of the RPE generated in the system with punctate state representation to the system with punctate action representation. Configurations are the same as those in Fig. 9, except that *κ* in this figure represents the degree of the effect of the TD RPE generated in the system with punctate state representation.

## Discussion

Referring to different streams of findings and suggestions in the literature of addiction and neuroscience of value learning, we have proposed a computational model of mechanisms that potentially contribute to the difficulty in resisting habitual behavior to obtain reward. In particular, in the model consisting of a series of state transitions towards the rewarded goal, we have shown that a sustained large positive RPE is generated upon goal reaching in the system with the goal-based reduced successor representation established under the Non-Resistant policy. We have further shown that the possible influence of DA/RPEs generated in the goal-based representation system on the system with punctate action representation through the spiral striatum-midbrain circuit could potentially enhance the propensity of non-resistant choice.

### Further possibilities about the effects of the generated RPEs on behavior

As shown in the Results, if the information of the RPEs generated in the system with the goal-based representation flows into the system with punctate action representation, it would act as an extra reward at the goal and thereby bias action values, potentially enhancing deterioration of the resistance to temptation. In the worse case, we speculate that the large positive RPEs upon goal reaching, coming from the system with the goal-based representation, could potentially even act as fictitious RPEs that cannot be fully canceled out by predictions within the punctate action representation system and thereby causes unbounded value increase and compulsion, similarly to what has been suggested for drug-induced DA (Redish, 2004). The anatomically suggested ventral-to-dorsal spiral influences (Haber *et al.*, 2000; Joel & Weiner, 2000) more precisely refer to the projections of more ventral parts of striatum to more dorsal parts of midbrain, rather than projections of more ventral parts of midbrain to more dorsal parts of striatum. Therefore, if every DA neuron in the dorsal parts of midbrain receives value information from both the ventral and dorsal parts of striatum, with *κ*: 1 − *κ* ratio as assumed in Fig. 8A, positive RPEs generated in the goal-based representation system can be canceled out by negative RPEs of action values at the level of inputs to the DA neuron. However, if there exist some DA neurons in the dorsal parts of midbrain that receive value information only from the ventral parts of striatum, such a cancelation cannot occur at the level of inputs. Then, if the amplitude of the positive RPE generated in the goal-based representation system is so large, resulting DA release from such DA neurons might not be able to be fully canceled out by a decrease or pause of DA release from surrounding DA neurons given the asymmetry of the positive and negative phasic responses of DA neurons (Bayer & Glimcher, 2005).

As we mentioned in the Results, our model assumes that the entire TD RPE flows from the ventral circuit to the dorsal circuit (Fig. 8A), different from (Keramati & Gutkin, 2013) but somewhat similar to (Takahashi *et al.*, 2008). If both the upcoming and previous values are sent via the direct striatum-midbrain connections, the suggested spiral connections could also convey both information, but it needs to be verified. If either of the upcoming and previous values (or both) is sent via the indirect pathway, spiral connectivity for both pathways is required for our model, and this also needs to be validated. Other than the possible effects of the spiraling RPE information, positive and negative RPEs generated in each system in our model themselves could cause subjective positive and negative feelings, respectively, given the suggestion that subjective momentary happiness of humans could be explained by reward expectations and RPEs (Rutledge *et al.*, 2014).

### Strengths of the present work/model

A strength of our model is that it does not assume drug-induced direct modulations of the DA system but still considers a key role of DA, and so our model can apply to any kinds of substance or non-substance reward and potentially explain the suggested similar involvements of the DA system in addictions to substance and non-substance rewards. Habitual, or even addictive, reward taking can arise not only for “DA-hijacking” substance but also for natural substance, such as food, or non-substance, such as gambling, gaming, smartphone use, or interaction/relation with other persons. Moreover, it has been suggested that the DA system is also involved in behavioral addiction to non-substance reward (Grant *et al.*, 2010). Specifically, there have been suggestions of possible relations of medicines of Parkinson disease to pathological gambling (Dodd *et al.*, 2005; Voon *et al.*, 2006) and of similar changes in the DA system in addiction to substance and non-substance such as game (Thalemann *et al.*, 2007) or internet (Hou *et al.*, 2012). In our model, resistance to temptation causes a large positive DA/RPE signal at the rewarded goal in the goal-based representation system. Crucially, different from the conventional DA/RPE response to reward, which disappears once the reward becomes predictable, the DA/RPE signal in the goal-based representation system continues to be generated. It has thus a similarity to the drug-induced DA release, providing a potential mechanism for the suggested similar involvements of the DA system in substance and non-substance addictions. Previous studies proposed mechanisms for, or applicable to, non-substance addiction related to state representation (Redish *et al.*, 2007), high DA release in the nucleus accumbens (Piray *et al.*, 2010), and the complexity of after-effects (Ognibene *et al.*, 2019). Our proposed mechanism is distinct from, and potentially complementary to, them.

Another, more general strength of the present work lies in its message that inaccurate value estimation due to low-dimensional state representation, and resulting sustained RPEs that could transmit from one system to another, can potentially lead to behavioral problems and even psychiatric disorders. The successor representation is a neurally implementable way of partially model-based RL, but one of its critical drawbacks is policy-dependence (Momennejad *et al.*, 2017; Russek *et al.*, 2017; Piray & Daw, 2019). Dimension reduction in state representation in the brain is generally suggested (Gershman & Niv, 2010; Niv, 2019), but it is inevitably accompanied by the risk of inaccuracy. The hierarchical cortico-basal ganglia structure has been suggested to have functional significances (Haruno & Kawato, 2006; Botvinick *et al.*, 2009; Frank & Badre, 2012; Collins & Frank, 2013; Baladron & Hamker, 2020), but it could relate to drug addiction (Keramati & Gutkin, 2013). The present work proposes that a combination of these negative sides can be related to behavioral problems in general, and to the difficulty in the cessation of undesired habits in particular.

### Drawbacks/limitations of the present work/model

The present model proposes a possible mechanism for why cessation of habitual reward-obtaining behavior is difficult, but does not explain why habitual or addictive reward-obtaining is initially shaped. In this regard, the present model is complementary to previous models for addiction. Also, although our model generally points to the empirically suggested similar involvements of the DA system in both substance and non-substance addiction, the results of our simulations do not specifically link to known behavioral or physiological results reported for addiction. For this, we will discuss possible neuroimaging experiments in the next section.

Next, our model critically depends on the assumption that the goal-based reduced successor representation is used in humans and implemented in the brain, but we could not find any direct evidence for them. As for behavioral evidence, we will discuss possible experimental validation in the next section. Regarding neural implementation, we found potentially supporting findings in the literature. Specifically, a finding that the BOLD signal in the ventromedial prefrontal cortex and hippocampus was negatively correlated with the distance to the goal in a navigation task (Balaguer *et al.*, 2016) appears to be in line with such a goal-based representation; if those regions engaged predominantly in the genuine successor representation in that task, their overall activity may not show a monotonic increase towards the goal. It is conceivable that the genuine successor representation can be encoded in the hippocampus (Stachenfeld *et al.*, 2017), but the reduced goal-based representation can become dominant through intensive training on a particular task or through long-standing habitual behavior towards a particular goal. Another study (Howard *et al.*, 2014) has shown that the BOLD signal in the posterior hippocampus was positively correlated with the path distance to the goal (increased as the path became farther) during travel periods whereas it was negatively correlated with an interaction between the distance and direction to the goal (increased as the path became closer and more direct) at decision points (and prior studies potentially in line with either of these results are cited therein (Spiers & Maguire, 2007; Morgan *et al.*, 2011; Viard *et al.*, 2011; Sherrill *et al.*, 2013)). The goal-based representation that we assumed can potentially be in line with the activity at decision points, rather than during travel periods, in that study.

Yet another important limitation of the present work is that we modeled the person’s resistance to temptation by directly setting the probability of “No-Go” choice rather than describing the mechanism of action selection (decision making) of the person who has an intention to quit the habitual reward-obtaining. In terms of value-based action selection, the Non-Resistant policy in our model is just optimal, and the Resistant policy is not. For this issue, we consider that in addition to the systems for value learning and value-based action selection / decision making, there would also exist distinct system(s) for rule learning and rule-based decision making, presumably including prefrontal (especially anterior prefrontal / fronto-polar) cortical circuits (Miller & Cohen, 2001; Strange *et al.*, 2001; Sakai, 2008). Rule can be set both externally (e.g., by law, or by other person) or internally (as a self-control). Rule-based behavior could theoretically be also regarded as a sort of value-based behavior, driven by punishments (negative rewards) given when breaking the rules, but would be more absolute or compulsory. Incorporation of the rule-based system into the model is an important future direction.

### Possible experimental validation, and clinical implication

The goal-based reduced successor representation, the critical assumption of our model, can be considered to be an example of reduced successor representation where each state is represented by the discounted future occupancies of not all the states but only the states with immediate rewards or punishments; such states themselves could become specifically represented through salience signals. It would be possible to conduct behavioral experiments to examine whether humans adopt such reduced successor representation or the genuine successor representation, somewhat similar to the experiments (Momennejad *et al.*, 2017) that compared the reevaluation of reward, transition, and policy. Specifically, if the dimension-reduced representation based on the states with immediate rewards/punishments is used, adapting to changes in reward placement (i.e., in what states reward is obtained) should be more difficult than adapting to changes in reward size.

At the neural/brain level, our model predicts that distinct patterns of RPEs are generated at each state leading to the rewarded goal in the systems with the goal-based representation (Fig. 3) and punctate state representation (Fig. 6) or genuine successor representation (Fig. 7), which could be encoded by DA released in different parts of the striatum. This prediction can potentially be tested by fMRI experiments and model-based analyses (O’Doherty *et al.*, 2007; Daw, 2011).

From clinical perspectives, it is essential to know whether the phenomena described by the present model actually occur in people who are trying to resist long-standing behavior to obtain reward, and whether the generated RPEs indeed contribute to the difficulty in cessation of such behavior. A potential way is to conduct brain imaging for those people executing a task that simulates their daily struggles against reward-obtaining behavior, including failures to resist temptation. If it is then suggested that the large positive RPE upon goal reaching generated in the system with the goal-based representation is an important cause of the difficulty, a possible intervention that is potentially effective is to provide alternative reward (physical, social, or internal) upon “No-Go” decisions until the state representation changes and approaches the one under the Resistant policy.

## Materials & Methods

Equations and parameters used are described in the Results. As described there, we set the number of states (*n*) to 10, and the time discount factor (*γ*) was varied over 0.97 ± 0.02, resulting in that the value at the start state was 0.95^9^ (≈ 0.63), 0.97^9^ (≈ 0.76), or 0.99^9^ (≈ 0.91) times of the value at the goal. We assumed 10 states because it seems intuitively reasonable to assume that the long-standing daily behavior to obtain a particular reward, such as going to a favorite pub for a beer after work, consists of around several to 10 distinct actions, e.g., clean the desktop, wear the jacket, wait for and get on the elevator, walk to the subway station, wait for and get on a train, walk to the pub, call the waitstaff, and order the beer. These series of actions would typically take dozens to tens of minutes. Given this, we determined the abovementioned range of time discount factor in reference to a study (Buono *et al.*, 2017), which examined temporal discounting for video gaming and found that the subjective value of video gaming 1 hour later was on average around 0.65 ~ 0.8 times of the value of immediate video gaming. Notably, however, the temporal discounting reported in that study appears to have near flat tails, indicating that it would not be well approximated by exponential functions, whereas we assumed exponential discounting.

In order to examine average behavior of the model across simulations, simulations were conducted 100 times for each condition. Among the 100 simulations, there were likely to be simulations, where “No-Go” choice was not taken at some state(s) at some episode(s). Such simulations, different from case to case, were not included in the calculations of the average and standard deviation of RPEs across simulations. There were also likely to be simulations, where “No-Go” choice was taken more than once at some state(s) at some episode(s). In such cases, generated RPEs were first averaged within an episode, and that value (i.e., a single value for each simulation) was used for the calculations of the average and standard deviation of RPEs across simulations. Simulations and figure drawing were conducted by using Python and R, respectively.

## Acknowledgements

K.M. is supported by Grant-in-Aid for Scientific Research (No. 20H05049) of the Ministry of Education, Culture, Sports, Science and Technology in Japan (http://www.mext.go.jp/en/). A.K. is supported by Grant-in-Aid for JSPS Fellows (No. 19J12156) of the Japan Society for the Promotion of Science (https://www.jsps.go.jp/english/). The authors thank Mr. Hirokazu Hatta for literature search on computational models of addiction.

## Data availability statement

Program codes for generating all the data presented in the figures are available in the GitHub (https://github.com/Kshimod/Reduced_SR_RL).

## Author contributions

K.M., K.S., and A.K. conceptualized the study. K.S. conducted the simulations and prepared the graphs. A.K. and K.M. validated the simulations and the graphs. K.M. supervised the project and prepared the original draft. K.S., A.K., and K.M. revised the draft. K.S. and A.K. contributed equally to this work.

